# Expression of substance P, NPY and their Receptors Is Altered in Major Depression

**DOI:** 10.1101/2022.12.14.516867

**Authors:** Swapnali Barde, Julio Aguila, Wen Zhong, Anna Solarz, Irene Mei, Josee Prud’homme, Miklos Palkovits, Gustavo Turecki, Jan Mulder, Mathias Uhlén, Corina Nagy, Naguib Mechawar, Eva Hedlund, Tomas Hökfelt

## Abstract

**BACKGROUND:** Major depressive disorder (MDD) is a serious disease and a burden to patients, families and society. Rodent experiments and human studies suggest that several neuropeptide systems, including substance P(SP)/tachykinin, neuropeptide Y(NPY) and their G protein-coupled receptors are involved in mood regulation.

**METHODS:** We assessed the transcript levels (qPCR) of SP/tachykinin and NPY systems in five regions from postmortem brains of male and female depressed subjects who committed suicide (DSS) and controls: dorsolateral prefrontal cortex (DLPFC), anterior cingulate cortex (ACC), the dorsal raphe nucleus (DRN), locus coeruleus (LC) and medullary raphe nuclei (MRN). We also analysed human LC neurons isolated using LCM with Smart-seq2 RNA sequencing.

**RESULTS:** Transcripts for all nine members were detected in male and female controls with marked regional variations of the raw CT values and with the highest levels for several tachykinin and tachykinin receptor transcripts in the DRN and for *NPY* and *NPYR* transcripts in the PFC regions. Significant sex differences for controls were recorded only in the DRN (*NPYR2* >in females) *and* LC (*TAC3* and *NPY* >in females). Elevated expression in DSS was recorded in (i) DLPFC for *SP, TAC* and *TAC3* in females, *SP* in males, and *NPYR1* in both sexes; and (ii) LC for all tachykinin family transcripts in females, *SP, TACR1* and *TACR3* in males, *NPY* in both sexes, and *NPYR1* in males.

**CONCLUSIONS:** The selective perturbation of neuropeptide systems in MDD patients may assist in the search for novel treatment strategies for subjects afflicted by this grave disorder.

## INTRODUCTION

Major depressive disorder (MDD) is a serious disease, afflicting more women than men and associated with much suffering and major costs for society (Wittchen, 2012; Murray et al., 2013; World Health Organization, 2017), and up till recently often also with treatment resistance (Akil et al., 2018). Interaction between genetic and environmental factors with stressful life events may represent important predisposing features (Nestler et al., 2002). Over decades, monoamine signaling has been implicated in the processes that lead to the symptoms of MDD, a view supported by the antidepressant effects of drugs that enhance serotonin and norepinephrine (NE) signaling, or both (Mongeau et al., 1997; Millan, 2006). More recently a paradigm shift, from monoamines to amino acids, has been announced based on breakthrough research, showing that the glutamate receptor antagonist ketamine has a rapid effect on treatment-resistant depression (Kadriu et al., 2019; Krystal et al., 2019). Also neuropeptides, the most diverse group of transmitters (Burbach, 2010), have been considered (Maubach et al., 1999; Hokfelt et al., 2003; Holmes et al., 2003; Nemeroff and Vale, 2005; Griebel and Holsboer, 2012; Hoyer and Bartfai, 2012).

Substance P (SP), neuropeptide Y (NPY) and their multiple receptors are widely expressed and distributed in the rodent and human brain, as shown in a large number of studies using various methods. Both systems have been analysed in extensive animal experiments regarding a possible role in various behaviors and pathological states, including mood derangements. Moreover, attempts have been made to explore a role and therapeutic potential of these peptide systems in human mood disorders. Thus, it was reported that a substance P (NK1) antagonist, MK-869 (Aprepitant, EMEND), has robust antidepressant activity in MDD subjects (Kramer et al., 1998). However, a phase 3 trial failed to reproduce the antidepressant effects of MK-869 (Keller et al., 2006), but this compound entered the clinic for treatment of chemotherapy-induced nausea (Hargreaves et al., 2011). Nevertheless, NK1 antagonists have been further discussed (Rupniak and Kramer, 2017) and explored in different contexts like, e.g., posttraumatic stress disorder (Frick et al., 2016), social phobia (Furmark et al., 2005), panic disorder (Fujimura et al., 2009) and alcoholism (George et al., 2008). Regarding NPY and treatment of depression in humans, a randomized controlled trial of intranasal administration of NPY to subjects with MDD has been performed (Mathé et al., 2020).

We have previously studied the expression of galanin and its three receptors, GALR1-3, in five regions of postmortem brains from depressed subjects who committed suicide (DSS) and controls: dorsolateral prefrontal cortex (DLPFC), anterior cingulate cortex (ACC), the dorsal raphe nucleus (DRN), locus coeruleus (LC)] and medullary raphe nuclei (MRN) (Barde et al., 2016); and we reported significant changes in transcript levels with opposite changes in DNA methylation (Barde et al., 2016). We have now processed the *same* tissue samples (albeit a reduced number, i.e. 168 vs 212 due to low yield for some), using qPCR analysis and with a focus on the SP/tachykinin and NPY systems. We show partly similar and parallel, but more limited changes compared to those seen for the galanin system, especially in the LC, a nucleus that recently has been thoroughly re-evaluated (Poe et al., 2020). We have therefore included RNA sequencing analysis of neurons isolated from the LC of 20 individuals. Finally, for comparison we will include some published results on the galanin system in the discussion (Barde et al., 2016). Part of these data were reported at a Nobel symposium in June 2022 (Barde et al., 2022).

## MATERIALS AND METHODS

### Brain samples

Postmortem brain tissue for qPCR analysis was obtained from the Douglas-Bell Canada Brain Bank. A total of 212 punched brain samples from five different regions were included (***Table 1***). Ethical approval for this study was obtained from The Institutional Review Board of the DMHUI, with written informed consent from the families, and from Karolinska Institutet (*SI Appendix, Material and Methods, Brain samples and Table S1)*. Fresh frozen post mortem brain tissue for RNA sequencing was obtained from the Netherland’s Brain Bank (NBB) and NIH Neurobiobank. LC tissue samples from a total of 20 individuals were included. Ethical approval for this work was obtained from the regional ethical review board of Stockholm, Sweden (EPN Dnr 2012/111-31/1). The work with human tissues was carried out according to the Code of Ethics of the World Medical Association (Declaration of Helsinki).

### RNA isolation and integrity analysis

Total RNA was isolated using RNeasy Plus mini kit (Qiagen). RNA quantity and quality were determined spectrophotometrically by using a ND1000 nanodrop (Saveen Werner). RNA integrity (RIN) was checked using Experion automated electrophoresis system (Bio-Rad). Samples that had RIN values of ≥4 were included in the analysis and the ones with very low concentrations/yield and RIN values ≤4 were excluded from the RT-qPCR analysis (i.e. n=168 punched samples were analysed).Total RNA was reverse transcribed to generate cDNA using High Capacity reverse transcription kit (Life Technologies) as per the manufacturer’s instructions. (*SI Appendix, Material and Methods)*.

### Quantitative real time PCR (qPCR)

RT-qPCR was performed as described previously (Le Maitre et al., 2013) with some modifications as described in detail in the Supplementary Information (SI). Expression stability of the endogenous controls across the five regions was previously checked (Barde et al., 2016). An NTC (no-template control) reaction and a RT-ve control reaction were used to check on for unspecific amplification and amplification from gDNA respectively. Relative fold changes were calculated by using the comparative CT method (2^-ΔΔCT^). (*SI Appendix, Material and Methods)*.

### Statistical Analysis

Statistical analysis was performed with GraphPad Prism 6 (GraphPad software, CA) StatView (SAS Institute Inc.). The different variables: RIN, age, PMI and tissue pH values between controls and suicides for the five brain regions were analyzed by multivariate ANOVA and two-tailed t-test for independent groups. Shapiro-Wilks test was used to test for Gaussian distribution. Significant outliers for qPCR fold change were analyzed by ROUT method and excluded if p<0.05. qPCR fold change data was analyzed by using the nonparametric Mann-Whitney U-test. *p*-values below 0.05 were considered significant, and *p*-values between 0.05 and 0.1 were considered to represent a trend.

### Laser capture microdissection, RNA sequencing and mapping of LC neurons

Frozen tissue samples originating from human pons and containing LC neurons were sectioned coronally at 10 um thickness using a cryostat. Sections were placed on precooled PEN membrane glass slides (Zeiss). Laser capture microdissection followed by Smart-seq2 RNA sequencing (LCM-seq) was carried out as described previously (Nichterwitz et al., 2016; Nichterwitz et al., 2018). LC neurons were selected based on their precise location and the presence of neuromelanin. We collected 50-120 neurons from each tissue sample and pooled these cells to generate one cDNA library of purified LC neurons/tissue sample for sequencing. Samples were sequenced using either the HiSeq2500 or the NovaSeq platforms (43 or 50 bp read length). The LC fastq files were mapped to the hg38 genome using STAR. Expression levels were determined using the rpkmforgenes.py software (http://sandberg.cmb.ki.se/rnaseq) with the Ensembl gene annotation. Counts were DESeq2 (v.1.34.0) normalized and log2 transformed.

## RESULTS

### Cohort demographics

There were no significant differences (mean ± S.D) between DSS and their matched controls in age (DSS vs. controls: 51.6 ± 15.4 years vs. 59.6 ± 15.4 years, *P*=0.08); postmortem interval (DSS vs. controls: 43.56 ± 24.16 hours vs. 44.17 ± 29.93, *P*=0.92); brain pH (DSS vs. controls: 6.65 ± 0.28 vs. 6.48 ± 0.34; *P*=0.12) or RIN (DSS vs. controls: 6.73 ± 1.67 vs. 6.54 ± 1.74; *P*=0.42). Demographic characteristic details of DSS and controls for each of the regions analyzed are provided in ***Table 1a***, and detailed information on each individual subject is provided in ***Table S1***. Of the 20 subjects used for RNA sequencing analysis, 30% were females (n=6) and 70% were males (n=14). The mean (± SD) age was 53.05 ± 15.4 years and postmortem intervals was 9.4 ± 6.19 hours, respectively. Details of these subjects are provided in ***Table 1b*** and ***Table S2***.

**Table 1a.** Demographic characteristics of the cohort used for qPCR.

|  | ACC 8/9 | ACC 24 | DRN | LC | MRN |
| --- | --- | --- | --- | --- | --- |
| **Sample size** | **38** | **35** | **35** | **32** | **28** |
| **Sex: %male controls (n)** | 26.3% (10) | 25.7% (10) | 25.72% (9) | 31.25% (10) | 25% (7) |
| **% male suicide (n)** | 26.3% (10) | 20% (7) | 28.57% (10) | 21.88% (7) | 28.6% (8) |
| **Sex: % female (n)** | 23.7% (10) | 28.6% (10) | 22.86% (8) | 25% (8) | 32.1% (9) |
| **% female (n)** | 23.7% (10) | 28.6% (10) | 22.86% (8) | 21.88% (7) | 14.3% (4) |
| **Age: Mean years ± SD** | 61.08±14.49 | 62.64±13.57 | 58.26±14.96 | 56.03±17.95 | 60.68±15.32 |
| **Postmortem interval: mean hours ± SD** | 43.96±29.41 | 49.31±28.88 | 45.07±28.89 | 45.78±29.53 | 45.48±28.39 |
| **pH value: Mean ± S.D.** | 6.51±0.35 | 6.57±0.35 | 6.52±0.37 | 6.48±0.31 | 6.52±0.40 |
| **RIN value: Mean±S.D.** | 7.02±1.23 | 6.82±0.46 | 6.02±1.26 | 7.44±1.12 | 6.68±0.86 |

**Table 1b.** Demographic characteristics of the cohort used for LCM-seq.

|  | LC |
| --- | --- |
| **Sample size** | **20** |
| **Sex: %male samples** | 70% (14) |
| **%female samples** | 30% (6) |
| **Age: Mean years ± SD** | 53.05±15.4 |
| **Postmortem interval: mean hours ± SD** | 9.4±6.19 |

### The effects of medication

Autopsy analysis is associated with an inherent problem of assessing the effect of medication, especially important when considering transcript levels and expression patterns. In fact, it has been described that, e.g., an antidepressant can exert epigenetic changes (Zimmermann et al., 2012; Gassen et al., 2015). Details of the analysis is described in SI. Based on this analysis, we conclude that in our study the various psychiatric medications show no significant effect on the gene expression of TAC and NPY systems in the five regions analyzed.

### qPCR study: the tachykinin system

#### Control subjects

*SP, TAC1* and their receptors (*TACR1, TACR2* and *TACR3*) were analysed (***Table S3***). The highest transcript levels were found for *TAC1* in DRN (rCt ∼20) followed by LC (rCT ∼24, i.e. 8-fold lower), MRN (rCT ∼25, i.e. 32-fold lower) and PFC and ACC (rCT ∼27 for both regions, i.e. 64-fold lower). *SP* exhibited a similar profile, although DRN levels were 4-fold lower (rCT ∼22) and PFC levels 2-fold (rCT ∼25) higher than *TAC1*, thus, the ranking order for TAC1 and SP is DRN>LC>MRN>PFC=ACC. Regarding receptors *TACR1* and *TACR3* showed the highest levels in DRN (rCt ∼26-27) versus PFC and ACC (∼31), whereas in MRN *TACR3* was 2-fold higher than *TACR1* (rCt ∼29 vs ∼31), followed by *TACR1* and *TACR2*. Thus, for TACR1 the order is DRN>LC>MRN>PFC=ACC, for TACR2: DRN>LC=PFC=ACC>MRN and for TACR3: DRN>LC>MRN>PFC>ACC (***Fig 1, Table S3***). In general, no differences in transcript levels were observed between male and females, except for LC: (i) whereas females have higher levels of *SP* mRNA (rCT ∼22 vs ∼24; i.e.4-fold), no difference was observed for *TAC*; (ii) males have higher transcript levels for *TACR1* (rCt ∼26 vs 28; 4-fold) and *TACR2* (rCT ∼30 vs 33; 8-fold) but lower levels of *TACR3* (rCT ∼30 vs 28; 4-fold) (***Fig 1, Table S3***).

**Fig 1.**
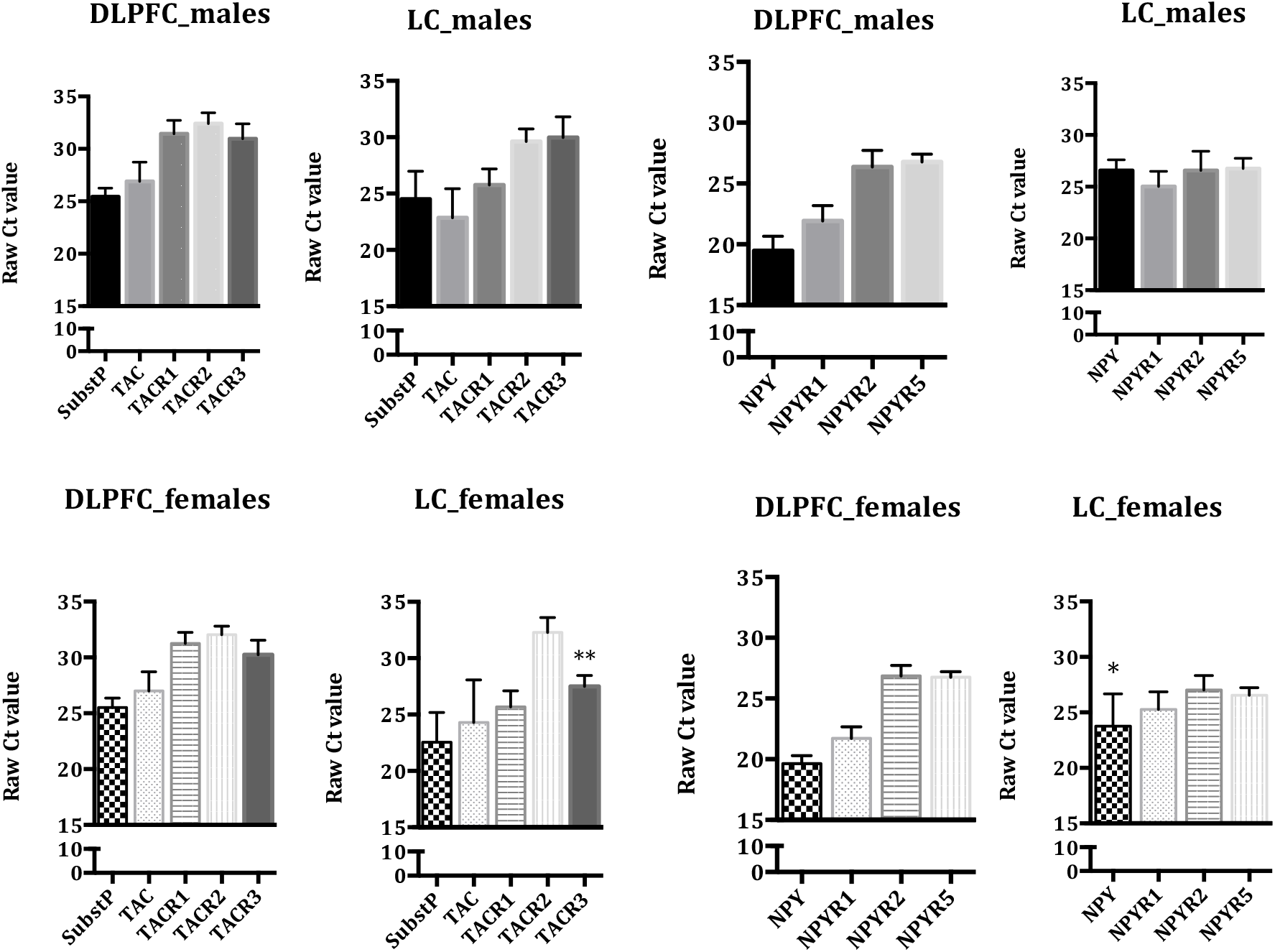
Raw Ct values in the DLPFC and LC of male and female controls for Substance P, TAC and its receptors (R1, R2 and R3). Threshold cycle number when the signal was observed is expressed as Raw Ct values in male and female control subjects for Substance P, NPY and their respective receptors. Lower Ct value corresponds to a more abundant transcript and vice-versa. Error bars represent mean ± S.E.M., n=8-10 per group, *P<0.05, **P<0.01.

The distribution of the *SP* system is well documented in the rat brain, in numerous biochemical and histochemical studies, reporting on the distribution of the peptide/protein and transcripts of substance P (Brownstein et al., 1976; Cuello and Kanazawa, 1978; Ljungdahl et al., 1978; Warden and Young, 1988) and of neurokinin A and B (Arai and Emson, 1986; Marksteiner et al., 1992; Mileusnic et al., 1999a).

Many studies have described the distribution of the substance P/neurokinin system in the human brain (e.g. (Ghatei et al., 1984; Bouras et al., 1986; Mai et al., 1986; Hornung et al., 1992; Mileusnic et al., 1999a). Detailed studies have been published on the DRN and LC (Halliday et al., 1990; Baker et al., 1991; Fodor et al., 1992), whereby there are in the DRN different SP neuron populations, some expressing serotonin (Halliday et al., 1990; Baker et al., 1991; Sergeyev et al., 1999).

The NK1-3 receptors have been studied in several species, especially in rats using immunohistochemistry (Saffroy et al., 1988; Mantyh et al., 1989; Lucas et al., 1992; Maeno et al., 1993; Nakaya et al., 1994; Bremner et al., 1996a; Mileusnic et al., 1999b). Autoradiographic analysis has shown NK1-3/SP binding sites/transcripts in the brain of rodents (Dam et al., 1988; Rigby et al., 2005), monkey (Nagano et al., 2006), and human (Jordan et al., 1995; Kus et al., 1998; Bensaid et al., 2001; Caberlotto et al., 2003; Rigby et al., 2005). Our results are in agreement with the findings in these studies. The *human* brain tachykinin system, especially the NK1 receptor, has been detected in particular in various forebrain areas of both men and women using positron emission tomography (PET) (Hargreaves, 2002; Bergström et al., 2004; Hietala et al., 2005; Okumura et al., 2008; Engman et al., 2012; Nyman et al., 2019). Finally, many RNAseq and single-cell studies present information on the type of cortical neurons expressing tachykinin family transcripts (Hodge et al., 2019; Mathys et al., 2019; Nagy et al., 2020; Sjöstedt et al., 2020; Zhong et al., 2022).

Taken together, our results support presence of moderate SP/TAC1 and TACR1-3 expression in different cortical regions with much higher transcript levels in DRN and LC of the male and female human brain, and here we also detect TACR1-3 transcripts, higher in DRN than in LC

#### Depressed subjects

In the DLPFC, *SP* mRNA levels were significantly increased in male and female DSS (P <0.05) when compared to controls (***Fig 2, Table 2***). *TAC1* mRNA levels were significantly increased only in female DSS (P<0.01). Similarly, *TACR2* and *TACR3* mRNA levels were significantly increased only in female DSS (P<0.05), implying distinct sex differences. *TACR1* transcript levels were unchanged (***Fig 2, Table 2***). In the ACC and MRN no significant changes were found in any of the five TAC family markers (***Table 2***).

**Table 2.** Overview of mRNA changes in male and female suicide subjects compared to controls.

|  | DLPFC |  | ACC |  | DRN |  | LC |  | MRN |  |
| --- | --- | --- | --- | --- | --- | --- | --- | --- | --- | --- |
|  | M | F | M | F | M | F | M | F | M | F |
| **SubsP** | ↑ | ↑ | - | - | ↑ | - | ↑ | ↑ | - | - |
| **TAC** | - | ↑↑ | - | - | - | - | - | ↑ | - | - |
| **TACR1** | - | - | - | - | - | - | ↑ | ↑ | - | - |
| **TACR2** | - | ↑ | - | - | - | - | - | ↑ | - | - |
| **TACR3** | - | ↑ | - | - | - | - | ↑ | ↑ | - | - |
| **NPY** | - | - | - | - | - | - | ↑↑ | ↑ | - | - |
| **NPYR1** | ↑ | ↑ | - | - | - | - | ↑ | - | - | - |
| **NPYR2** | - | - | - | - | - | - | - | - | - | - |
| **NPYR5** | - | - | - | - | - | - | - | - | - | - |
| **GAL** | ↓↓ | ↑ | - | - | ↑↑ | ↑↑ | ↑↑ | ↑↑ | ↑ | ↑ |
| **GALR1** | ↑ | ↑ | - | - | ↑ | - | ↑ | - | - | - |
| **GALR2** | - | - | - | - | - | - | - | - | ↓↓ | - |
| **GALR3** | ↓↓ | - | - | - | ↑ | ↑↑ | ↑ | ↑↑ | ↑ | ↑ |

The arrows represent statistical significance where upward arrow signifies increase in gene expression and vice-versa. One arrow represents $p < 0.05$ and two arrows represent $P < 0.01$ .
DLPFC, Dorsolateral Prefrontal Cortex; ACC, Anterior Cingulate Cortex; DRN, Dorsal Raphe Nucleus; LC, Locus Coeruleus; MRN, Medullary Raphe Nucleus.

**Fig 2.**
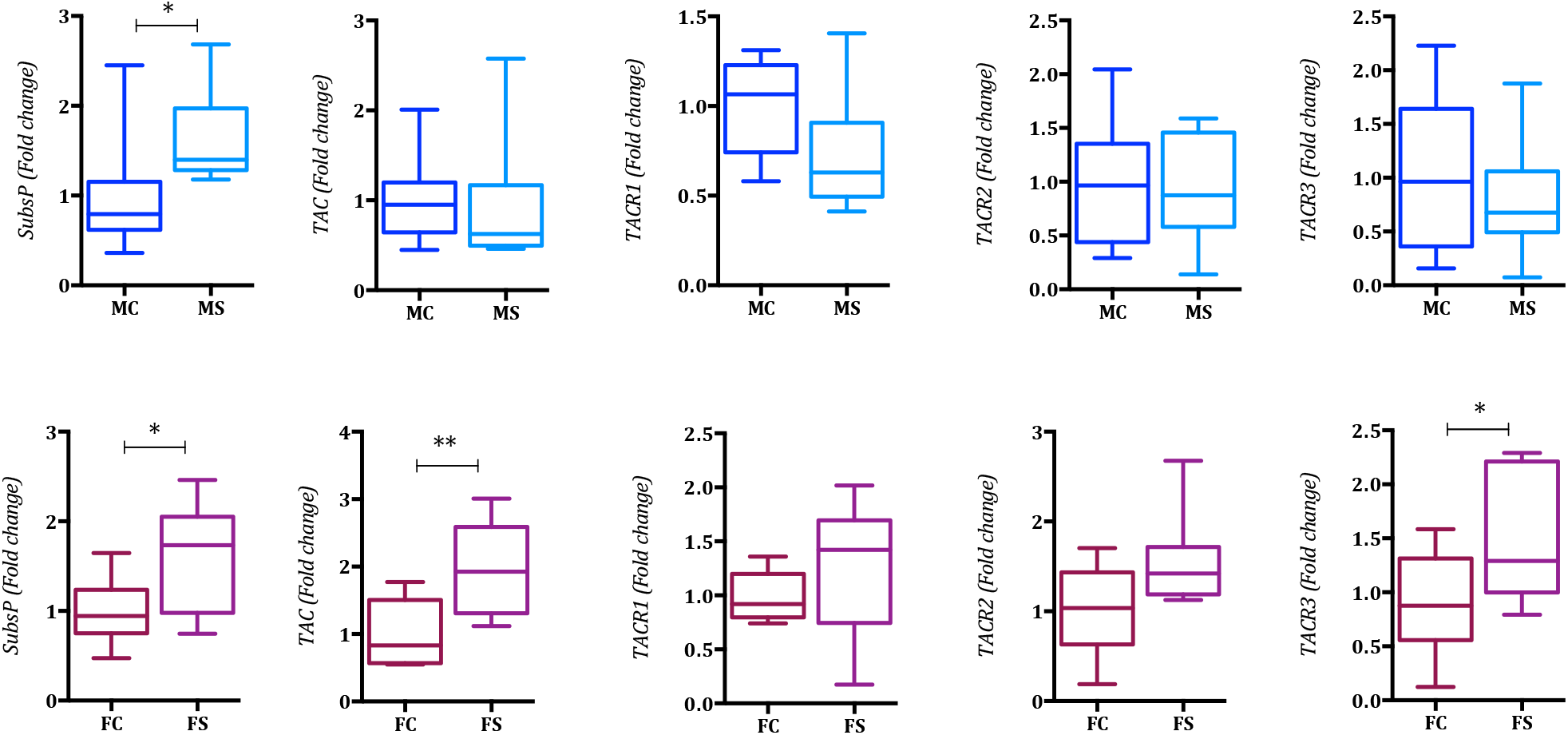
TAC and its respective receptors mRNA expression in the dorsolateral prefrontal cortex (BA 8/9) of male and female depressed suicide (DS) subjects. Differential RNA expression expressed as fold change in male (panel A) and female (panel B) depressed suicide subjects. Data are represented as fold change which is calculated using ddCt i.e normalized with endogenous transcript expression values and controls. Error bars represent mean ± S.E.M., n=10 per group, *P<0.05, **P<0.01.

Only *SP* mRNA levels were significantly elevated in the DRN of male DSS (P <0.05) but not of females (***Table 2***). In the LC, however, *SP* mRNA levels were significantly higher in male and female DSS (P <0.05) when compared to controls, and *TAC* transcript levels were significantly elevated only in female DSS (P <0.05). Regarding receptors, *TACR1* and *TACR3* mRNA levels were significantly higher in both male and female DSS (P<0.05), but *TACR2* levels only in female DSS (***Fig 3, Table 2***).

**Fig 3.**
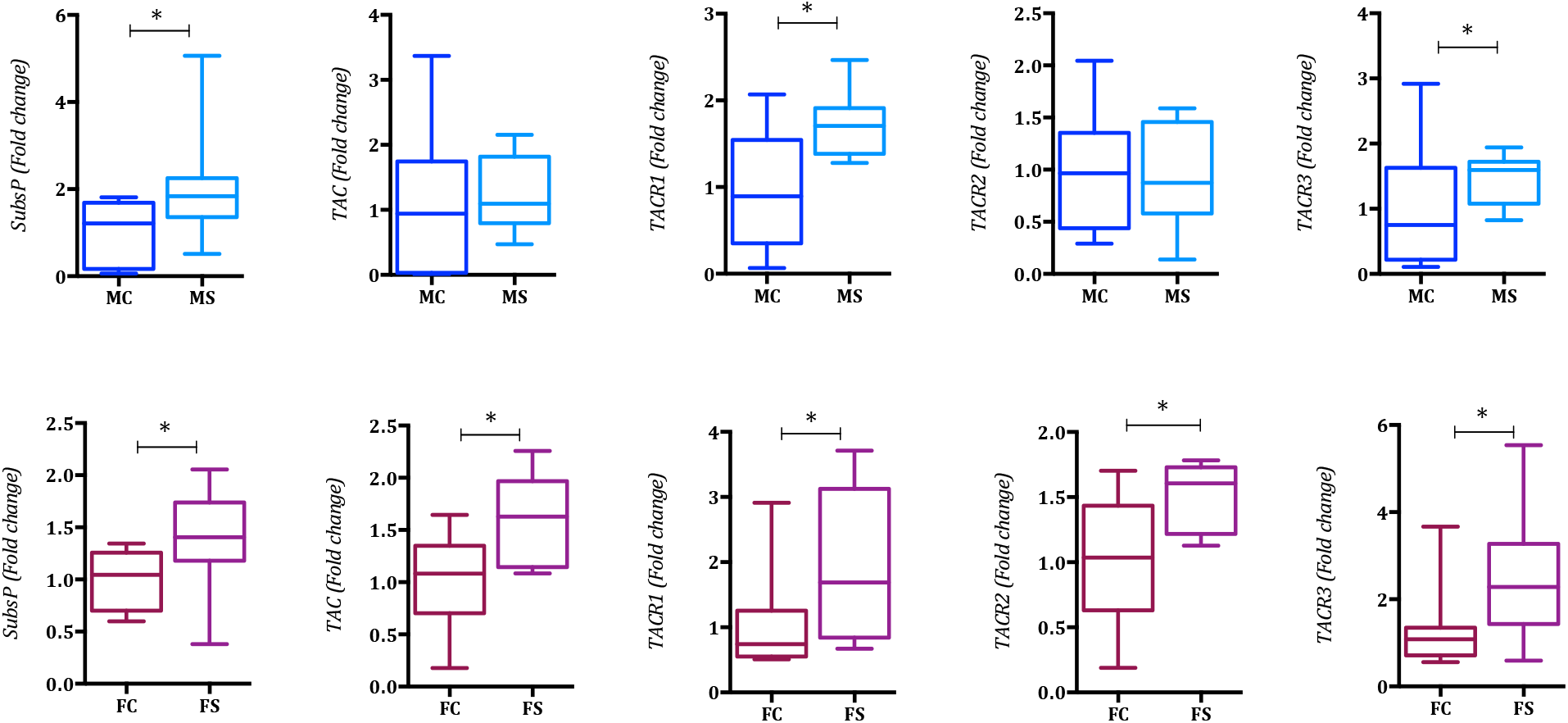
TAC and its respective receptors mRNA expression in the locus coeruleus (LC) of male and female depressed suicide (DS) subjects. Differential RNA expression expressed as fold change in male (panel A) and female (panel B) depressed suicide subjects. Error bars represent mean ± S.E.M., n=10 per group, *P<0.05, **P<0.01.

### qPCR study: the neuropeptide Y system

#### Control subjects

The *NPY* transcript levels were by far highest in the two cortical regions: the expression in DLPFC (rCt∼19) was ∼2-fold higher than in ACC (rCT∼20). In the brain stem regions, the highest expression was observed in the LC (rCt∼25) followed by DRN (rCT∼26), and then MRN (rCT ∼27). The ranking order is thus DLPFC>ACC>>>LC>DRN>MRN (***Fig 1, Table S4***). *NPYR1* was most abundant in the DLPFC (>ACC>LC>DRN>>MRN (rCT∼21>>23>>25>>27>>>30), whereas *NPYR2* levels were in general low: DLPFC (rCT∼27)=DRN>LC>ACC=MRN (rCT∼29). *NPYR5* was expressed at approximately similar levels in DLPFC, ACC, DRN and LC (rCT∼27), but levels were ∼16-fold lower in the MRN (rCT∼31). A sex difference was only observed in the LC: *NPY* transcript levels were ∼8-fold higher in females versus males (rCt 23.8 vs 26.6) (***Fig 1, Table S4***).

NPY is widely expressed in the rat brain (Chronwall et al., 1985; Yamazoe et al., 1985; de Quidt and Emson, 1986). This is true also for the human brain (Adrian et al., 1983; Dawbarn et al., 1984), including cerebral cortex (Chan-Palay et al., 1985; Van Reeth et al., 1987; Hornung et al., 1992; Mikkelsen et al., 1993). Studies on rat (Smith et al., 1994) and mouse (Fu et al., 2010) have revealed low numbers of NPY-positive neurons in the DRN, likely not monoaminergic ones. NPY has been detected in norepinephrine (NE) neurons in LC both in *rat* (Everitt et al., 1984; Sawchenko et al., 1985) and *human* (Hökfelt et al., 1983; Fodor et al., 1992). The distribution of the receptors has been described using multiple methods in *rat* (Giardino et al., 1989; Ohkubo et al., 1990; Widdowson, 1993; Caberlotto et al., 1997; Dumont et al., 2000; Kopp et al., 2002; Stanic et al., 2006) and *human* (Westlind-Danielsson et al., 1987; Martel et al., 1990; Widdowson, 1993; Caberlotto et al., 1997; Dumont et al., 2000).

PET studies on the *human* NPY system are rare, focusing on the Y1 receptor (Hostetler et al., 2011), even if ligands have been generated (Fonseca et al., 2022). Also for the NPY system there are many RNAseq and single-cell studies with focus on cortex (Hodge et al., 2019; Mathys et al., 2019; Nagy et al., 2020; Sjöstedt et al., 2020; Zhong et al., 2022).

#### Depressed subjects

*NPY* mRNA levels were significantly increased only in the LC, both of male and female DSS (P<0.01 and P<0.05 respectively, ***Fig 4, Table 2***). In the DLPFC only the *NPYR1* mRNA levels were significantly higher, both in male and female DSS (P<0.05 for both), and in the LC only of male DSS (P<0.05) (***Fig 4, Table 2***).

**Fig 4.**
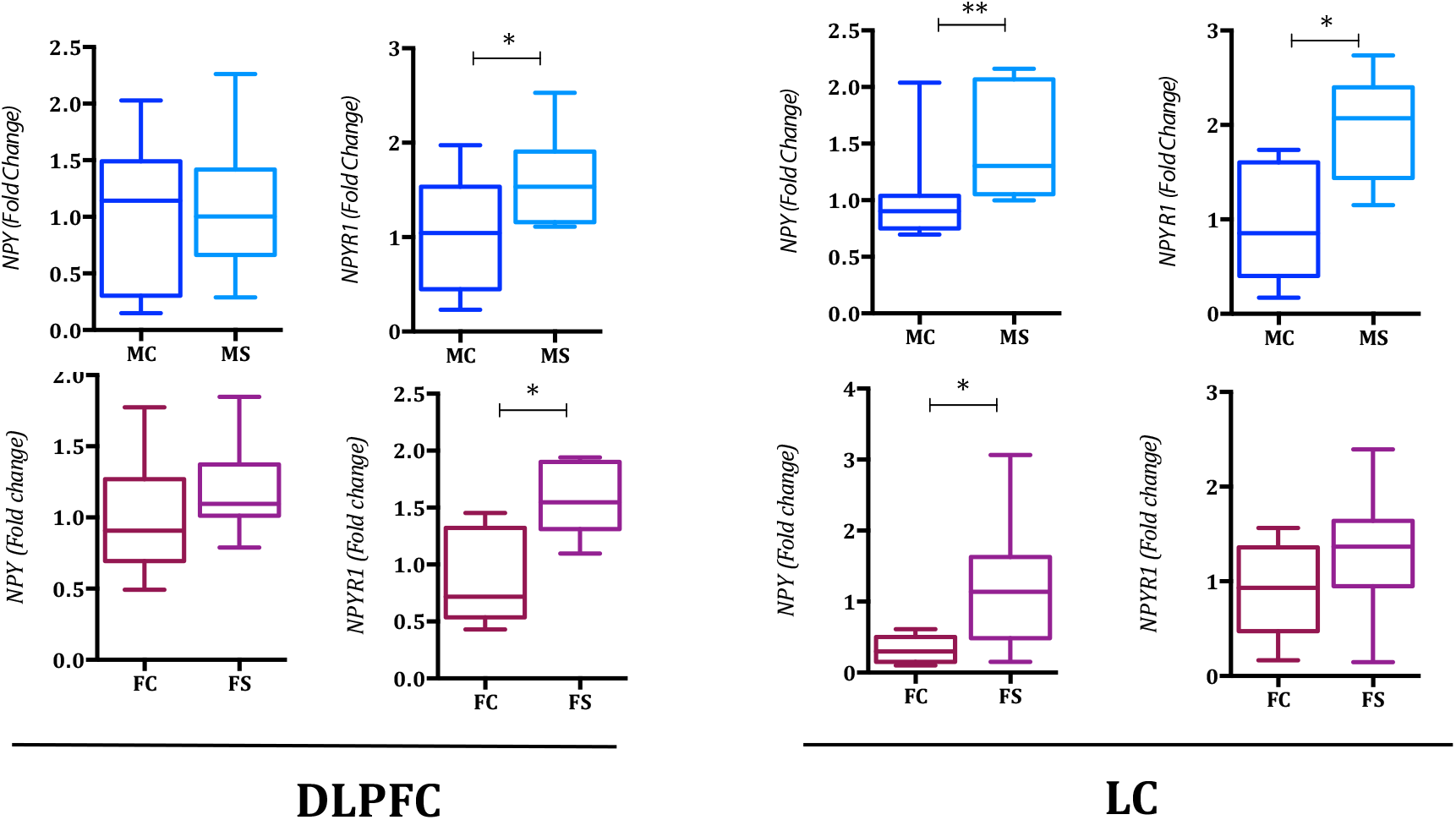
NPY and NPYR1 mRNA expression in dorsolateral prefrontal cortex (BA 8/9) and locus coeruleus (LC) of male and female depressed suicide (DS) subjects. Differential RNA expression expressed as fold change in male (panel A) and female (panel B) depressed suicide subjects. Error bars represent mean ± S.E.M., n=10 per group, *P<0.05, **P<0.01.

### LCM sequencing of LC neurons isolated from human post mortem tissues

Our study includes the tachykinin, NPY and galanin systems. Analysis of 20 samples using log2 transformation of normalized count values revealed that (i) TAC1 is present in 13 samples (mean: 4.24, SD: 4.01); TAC3 in 1 sample (mean: 0.10, SD: 0.46); TACR1 in all samples (mean: 7.18, SD: 1.34); TACR2 in 7 samples (mean: 0.97, SD: 1.88); TACR3 in in almost all, i.e. 19 samples (mean: 6.87, SD: 1.81). (ii) NPY is present in 10 samples (mean: 4.06, SD: 4.54); NPY1R in 4 samples (level: 0.63, SD: 1.64); NPY2R in 2 samples (mean: 0.39, SD: 1.24) and NPY5R in 1 sample (mean: 0.05). (iii) GAL is present in all 20 samples (mean: 5.88, SD: 2.2); GALR1 in 12 samples (mean: 2.22, SD: 2.27); GALR2 in 5 samples (mean: 0.38, SD: 1.04); GALR3 was not present in any of the 20 samples. See ***Fig 5***.

**Fig 5.**
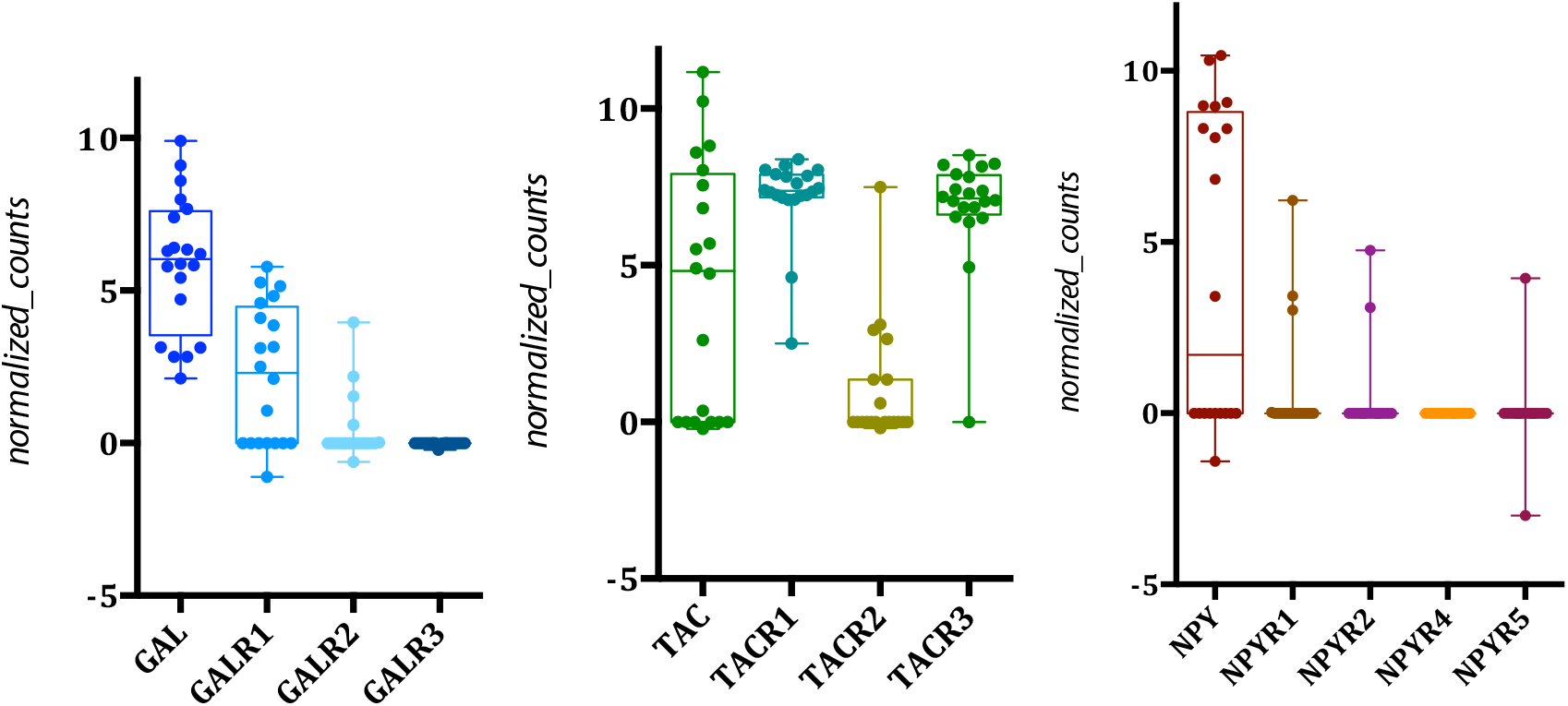
LCM-seq data for galanin, NPY and TAC family from isolated neurons of the human LC. RNA sequencing data analysed by log2 transformation of normalized data. Normalized counts for galanin and its receptors (A), TAC and its receptors (B) and NPY and its receptors (C) is shown.

Taken together there is a robust expression of TAC1, NPY, GAL, TACR1 and TACR3. Even if NPY was detected in fewer samples, two of the samples had a higher count than any of the GAL samples. The large variation in counts for the ligands is in agreement with an ISH study showing large differences in radioactive GAL signal between individual LC neurons (Le Maitre et al., 2013). The excitatory receptors (NK1 and NK3) are abundant, in all cells and fairly evenly expressed, as are the inhibitory receptors (GalR1 and NPYR1), even if in fewer cells. A surprise is perhaps that TAC1 is expressed in noradrenergic LC neurons.

## DISCUSSION

The present study provides novel, basic and quantitative information on the expression levels of five transcripts for the tachykinin and four for the NPY system in five regions of the male and female human postmortem brain, both from controls and DSS. Four of these regions have been associated with MDD. We show marked differences in expression levels of these two systems among the regions and some significant variance between sexes as well as several differences between controls and DSS. We include, in the discussion, for comparison data on the galanin system from a previous study (Barde et al., 2016) summarized in ***Table 2, Table S5***, that thus includes three peptide systems.

### Normal distribution patterns and sex differences

The present results confirm that the tachykinin system, in general, is strongly expressed in the lower brain stem, in LC and especially in DRN, where *TAC* and *SP* as well as *TACR1-3* transcript levels are higher than in any other region (*TAC*>*SP*>>*TACR1*=*TACR3*). However, *SP*/*TAC* levels are significant also in the two PFC regions (***Fig 1, Table S3***). Thus, the tachykinin system seems to be more strongly expressed in the human than in rat cortex (*cf*. (Ljungdahl et al., 1978)). In contrast, *NPY* is most highly expressed in the cortical regions, whereas the receptor levels are similar in the different regions. Sex differences for controls are only found in two regions: *TACR3 and NPY* are higher in the female LC, and in DRN *NPYR2* his higher in female DRN. The galanin peptide is most highly expressed in the LC sample (*LC*>>*DRN*>*ACC*=*DLPFC*) (Barde et al., 2016).

### In which neurons are the transcripts expressed and what are the potential effects of the ligands?

This is a bulk study, i.e. many neuron and non-neuronal populations are included in the dissected sample. It is therefore important to try to define, in which population(s) the studied transcripts likely are expressed. For PFC we therefore mainly use data from two single-cell/nuclei studies (Mathys et al., 2019; Nagy et al., 2020); for DRN and LC from Weber et al.(Weber et al., 2022) and from our own LCM-seq study on LC. Also, it is important to note that the results likely mainly reflect the sum of transcript levels in *cell bodies*, that could include more than one subpopulation of neurons and could range from a few neurons with high levels to many with low levels. This is clearly shown in the mentioned single-cell studies (Mathys et al., 2019; Weber et al., 2022).

In order to understand the mechanism of action of the three peptides, we include what type of electrophysiological effects that they may exert. However, whereas many systems have been histochemically identified in the *human* brain, very few electrophysiological experiments have been carried out on human brain samples. We will therefore refer to some relevant animal studies.

With regard to DLPFC both the transcripts for *TAC1* and *TAC3* and *NPY* are primarily localized in GABAergic neurons, but *NPY* also in glutamatergic neurons (Mathys et al., 2019; Nagy et al., 2020). *TACR1* is mainly expressed in GABAergic neurons, *TACR3* mainly in glutamatergic neurons, *NPY1R* in GABAergic and glutamatergic neurons and *NPY5R* in glutamatergic neurons (Mathys et al., 2019; Nagy et al., 2020).

In the rat forebrain SP is a powerful excitatory agent when applied to cortical neurons (Jones and Olpe, 1984), while NPY via postsynaptic Y1 receptors may cause inhibition, and via presynaptic YR2 receptors inhibition of glutamate release (Sperk et al., 2007). In electrophysiological studies on the *human* temporal lobe NPY strongly decreased stimulation-induced excitatory postsynaptic potentials via NPYR2 receptors (Ledri et al., 2015).

A recent single-nucleus study has reported on 5-HT neurons in the most caudal part of the human DRN (intermingling with NE-LC neurons) (Weber et al., 2022). The results show that the 5-HT neurons are characterized by (i) a robust expression of *TAC1* and a modest signal for *TACR1* and *TACR3*; (ii) in general very low expression of the NPY family members - only *NPYR1* is detected; and (iii) a weak expression of *GalR2* and *GALR3*, but no signal for *GAL* or *GALR1*. All transcripts are in addition expressed at various levels in excitatory and inhibitory neurons, as well as in other cell types (Weber et al., 2022).

In rat, intracellular recordings from 5-HT neurons show a robust increase in excitatory postsynaptic currents after administration of either SP (TACR1 agonist) or senktide (NKB; TACR3 agonist); however, this effect is not direct but is mediated via stimulation of, possibly local, glutamatergic neurons (Liu et al., 2002). Galanin has been reported to hyperpolarise most 5-HT-sensitive neurons (Xu et al., 1998). Sharkey et al. (Sharkey et al., 2008) instead found that galanin acts via GalR1 on GABA interneurons and via GalR2 on 5-HT neurons.

In rat, intracellular recordings from 5-HT neurons show a robust increase in excitatory postsynaptic currents after both administration of either substance P (TACR1 agonist) or senktide (NKB; TACR3 agonist); however, this effect is not direct but is mediated via stimulation of, possibly local, glutamatergic neurons (Liu et al., 2002). Galanin has been reported to hyperpolarise most 5-HT-sensitive neurons (Xu et al., 1998). Sharkey et al. (Sharkey et al., 2008) instead found that galanin acts via GalR1 on GABA interneurons and via GalR2 on 5-HT neurons.

In the human LC region, our analysis on isolated LC neurons shows that *TAC1, NPY* and GAL, are robustly expressed (*GAL*>*TAC1*=*NPY*) in NE-LC cells, as are *TACR1* and *TACR3*, and to a lesser extent *GALR1*. These results agree with the above mentioned single-nucleus paper (Weber et al., 2022). However, an unexpected finding is that *GalR1* is not only is expressed in NE-LC cells (at modest levels) but mainly and robustly in oligodendrocyte precursor cells (Weber et al., 2022). All transcripts are in addition expressed at various levels in excitatory and inhibitory neurons and often in other cell types (Weber et al., 2022).

In rodents, SP has a direct excitatory effect on NE-LC neurons (Guyenet and Aghajanian, 1977; Engberg et al., 1981; Cheeseman et al., 1983; Olpe and Steinmann, 1991). In contrast, NPY has a depressant effect (Olpe and Steinmann, 1991) and potentiates the NE-induced inhibition, via NPYR2 (Illes et al., 1993). Several studies have shown that galanin reduces firing of LC neurons (Seutin et al., 1989; Sevcik et al., 1993; Pieribone et al., 1995).

### Expression of the tachykinin and NPY (and galanin) system are changed in MDD

The present and a previous study suggest, as indicated in ***Table 2***, that the three neuropeptide systems (tachykinin, NPY and galanin) are related to MDD mainly in two regions: DLPFC and LC. More restricted changes were observed in DRN and MRN, both essentially associated with the galanin system. We included MRN, with projections to the spinal cord, as a control region likely not reacting in MDD. In fact, there were no changes in the tachykinin or NPY system. The changes in the galanin system therefore came as a surprise but will not be discussed here.

Regarding DLFPC in MDD *SP, TAC, TACR2, TACR3, NPYR1, GAL* and *GALR1* are upregulated in females, in males only *SP* and *GALR1*, possibly in agreement with the notion that women are more likely to be afflicted by MDD (Kessler et al., 2003). Interestingly, *GAL* and *GALR3* are robustly downregulated in males. Stockmeier et al. (Stockmeier et al., 2002) have reported decreased binding of [125I]BH-SP in orbitofrontal cortex (BA 47) in MDD versus our increased levels of *TACR2* and -*R3*. Regarding the NPY system in MDD, a recent study by Sharma et al. (Sharma et al., 2022) shows upregulation of *NPYR1, NPYR2* and decreased *NPY* transcript levels, whereas we only detected upregulation of *NPYR1*, however both in males and females. Previously Caberlotto and Hurd (Caberlotto and Hurd, 2001) have reported increased *NPYR2* levels in layer IV in DSS. Overall our own findings suggest elevated excitatory neurokinin signaling in DLPFC in MDD and confirm elevated inhibition via NPYR1 in MDD.

Regarding the galanin system in the human PFC, our RNAscope analysis shows that both the *GAL, GALR1* and -*R3* transcripts are mainly expressed in glutamatergic projection neurons (Zhong et al., 2022). We have hypothesized that galanin via the inhibitory GalR3 on intra-telencephalic pyramidal neurons protects against glutamatergic over-excitation; GALR1 in pyramidal neurons may instead influence thalamic signaling (see ***Fig S6***, (Zhong et al., 2022). The downregulation of GAL and GALR3 could thus attenuate this protective mechanism, alternatively indicating exhaustion of protection/resilience mechanisms. *SP, TAC* and *NPY* are, on the other hand and as said, mainly expressed in GABAergic interneurons (DeFelipe et al., 2013).

In the present study, LC is the most strongly affected region, with increased levels of several transcripts of the three peptide systems, both in males and females. The increase in ligand levels and receptor availability in MDD suggests a more intense signaling of the respective system. For example, the NA neurons may be activated by a parallel increase in SP release and increased availability of TACR1 receptors. This likely elevated excitation by the tachykinin system may be counteracted by the mainly inhibitory peptides, NPY and galanin. In fact, both NPY (Heilig, 2004; Morales-Medina et al., 2010; Cohen et al., 2012; Reichmann and Holzer, 2016; Sabban et al., 2016; Kautz et al., 2017) and galanin (Sciolino et al., 2015; Hökfelt et al., 2018) have at the level of the LC been suggested to be involved in resilience. We hypothesize that in MDD the excitatory, prodepressive drivers like tachykinins, corticotropin releasing hormone (CRH) and hypocretin/orexin have ‘won over’ anti-depressive, ‘resilience-inducing’ factors like NPY and galanin.

## CONCLUDING REMARKS

Taken together these findings in general support the importance of both the tachykinin (Maubach et al., 1999; Stout et al., 2001; Santarelli and Saxe, 2003; Ebner et al., 2009; Griebel and Holsboer, 2012; Schank and Heilig, 2017) and NPY (Kask et al., 2002; Redrobe et al., 2002; Heilig, 2004; Harro, 2006; Morales-Medina et al., 2010; Wu et al., 2011; Kautz et al., 2017; Nahvi and Sabban, 2020; Domin, 2021) systems in mood related disorders, and there is also evidence that the galanin system is involved (Unschuld et al., 2010; Juhasz et al., 2014; Barde et al., 2016; Gonda et al., 2018). Of these three systems galanin and its receptors are the molecules that are most strongly modulated in MDD. We believe that the changes in levels of transcripts reported here support the relevance of further efforts to develop novel antidepressants based on peptidergic signaling.

## LIMITATIONS

The present study only reports on transcripts, i.e. no information on peptides/proteins is included. However, we cite numerous reports based on radioimmunoassay or immunohistochemistry evidencing, in most cases, presence of the translational products. The present samples had been stored at -80° C for several years longer than those analysed in the study on the galanin system by Barde et al. (Barde et al., 2016). The fact that e.g. NPY mRNA levels (still) are very high in cortex speaks against impaired quality of tissue samples. Nevertheless, this aspect should be considered, when we compare results from that and the present study. We have in several cases speculated about the role of the peptide systems in MDD: further ISH analysis and immunohistochemistry (when receptor antibodies become available) would allow validation of such hypotheses. Finally, we here only deal with three peptide systems. For example, recent single-nucleus study has revealed expression of several further peptides in the human NE-LC neurons, and a multitude of neuropeptide-containing nerve endings possibly innervating the NE-LC neurons has been described.

## ACKNOWLEDGEMENTS AND DISCLOSURES

This project is presently supported by the Swedish Research Council (04X-2887), the Swedish Brain Foundation, The Arvid Carlsson Foundation and grants from Karolinska Institutet. Earlier phases of this project were supported by a NARSAD Distinguished Investigator Award (2009), the European Community (NewMood, LSHM-CT-2004-503474; 2004-2008), AFA Insurance (2008) and by a five-year Unrestricted Bristol-Myers-Squibb Neuroscience Grant. Of particular importance were generous grants over a longer period from the Marianne and Marcus Wallenberg Foundation (1998-2009), and from the Knut and Alice Wallenberg Foundation. SB was supported by a postdoctoral stipendium from Hjärnfonden (Swedish Brain Foundation) and from SSMF (Swedish Society for Medical Research). JA was supported by a postdoctoral stipendium from SSMF. Human post-mortem tissues for LCM experiments were kindly received from the Netherlands Brain Bank (NBB) (www.brainbank.nl) and NIH Neurobiobank (www.neurobiobank.nih.gov). RNA-seq experiments on LC were supported by funding from Parkinsonfonden (grants: 1413/2022 and 1328/21). WZ was supported by the SciLifeLab & Wallenberg Data Driven Life Science Program (grant: KAW 2020.0239). NM is a CIHR New Investigator and FRQ-S Chercheur-boursier. The Douglas-Bell Canada Brain Bank is supported by the *Réseau québécois sur le suicide, les troubles de l’humeur et les troubles associés* (FRQ-S) and by a PSG grant from Brain Canada. Support was provided by the MTA-SE-NAP B Genetic Brain Imaging Migraine Research Group, Hungarian Academy of Sciences, Semmelweis University (KTIA_NAP_13-2-2015-0001); by the National Development Agency (KTIA_NAP_13-1-2013-0001), Hungarian Brain Research Program (KTIA_13_NAP-A-II/14); and the Hungarian Academy of Sciences (MTA-SE Neuropsychopharmacology and Neurochemistry Research Group).

The authors declare no conflict of interest.

