## Supplementary material for "Expression of substance P, NPY and their Receptors Is Altered in Major Depression": SI Text

[1]Department of Neuroscience, Karolinska Institutet, Stockholm, Sweden; [2]Department of Biochemistry and Biophysics, Stockholm University, 106 91, Stockholm, Sweden; [3]Department of Cell and Molecular Biology, Karolinska Institutet, 171 77, Stockholm, Sweden; [4]Science for Life Laboratory, Department of Protein Science, KTH Royal Institute of Technology, Stockholm, 11428, Sweden; [5]McGill Group for Suicide Studies, Douglas Mental Health University Institute, Verdun, QC, Canada; [6]Department of Psychiatry, McGill University, Montréal, QC, Canada; [7]The Hungarian Academy of Sciences, Budapest, Hungary and Human Brain Tissue Bank and Laboratory, Semmelweis University, H-1085, Budapest, Hungary.

### SI Materials and Methods

**Brain samples.** Postmortem brain tissue was obtained in collaboration with the Quebec Coroner's Office and the Suicide section of the Douglas-Bell Canada Brain Bank (Douglas Mental Health University Institute, Montreal, Quebec, Canada). A total of 212 punched brain samples from five different regions of 61 individuals (controls and depressed suicides) were included (Table 1, Table S1). All individuals were of French-Canadian origin, a homogeneous Caucasian population matched for postmortem interval (PMI; interval between death and freezing of the brain), age and tissue pH. Psychological autopsies were performed for both cases and controls as described previously, and diagnoses were established by a panel of psychiatrists based on DSM-IV criteria. Subjects in the control group died suddenly from accidental or natural causes. Samples were obtained from five different regions: dorsolateral prefrontal cortex (BA 8/9), anterior cingulate cortex (BA 24), dorsal raphe nucleus (DRN), locus coeruleus (LC), and the medullary raphe nuclei (MRN).

Suicide and sudden-death control brains underwent a process known as a psychological autopsy to retrieve phenotypic information. This proxy-based interview procedure is the accepted standard to obtain diagnostic information postmortem. Briefly, a few months following death, families were contacted and the person best acquainted with the deceased was recruited to undergo a series of structured interviews. These interviews were supplemented with information from archival material obtained from hospitals, the Coroner's office and social services. Following the interviews, clinical vignettes were produced and assessed by a panel of clinicians to generate DSM-IV diagnostic criteria. As detailed elsewhere (Dumais et al., 2005), the psychological autopsies provide socio-demographic characteristics, social developmental history, DSM-IV axis I diagnostic information and behavioral traits.

We also obtain toxicological assessments, as well as complete information on medication prescription.

The dorsal raphe nucleus and locus coeruleus were dissected using the following coordinates from a human brainstem atlas (Paxinos, 1995) : dorsal raphe nucleus from Obex+32 to +39, locus coeruleus from Obex +24 to +31, and medullary raphe nuclei (obscurus and magnus raphe) from Obex 0 to +16). It should be noted that the samples labeled "locus coeruleus", and especially "dorsal raphe nucleus" include many different neuron populations that are not nordrenergic and serotonergic, respectively. Ethical approval for this study was obtained from The Institutional Review Board of the Douglas Mental Health University Institute, and written informed consent was obtained from the family of each deceased subject before inclusion in the study. The Karolinska Institutet group has a permission to process post mortem brain samples (Regional Ethical Board in Stockholm, No. 2013/474-31/2).

**RNA isolation and integrity analysis.** Total RNA from each of the 212 samples was isolated using RNeasy Plus mini kit (Qiagen). RNA quantity and quality were determined spectrophotometrically by using a ND1000 nanodrop (Saveen Werner). RNA integrity was checked using Experion automated electrophoresis system (Bio-Rad). All samples that showed and an RNA integrity number (RIN) higher than 5 was considered good quality RNA samples, and higher than 8 were considered as perfect (Fleige and Pfaffl, 2006). Samples with very low RNA concentrations and integrity (RIN <4) were excluded from the RT-qPCR analysis (n=168 were processed for TAC, NPY and their respective receptor transcripts). Total RNA was reverse transcribed to generate cDNA using High Capacity reverse transcription kit (Life Technologies) as per the manufacturer's instructions.

**Quantitative real time PCR (RT-qPCR).** RT-qPCR was performed as described previously (Barde et al., 2016) with some modifications. Briefly, 500 ng of RNA was reverse transcribed using the High capacity reverse transcription kit (Life Technologies) and the cDNA was subject to 40 cycles of amplification and by using TaqMan gene expression assays and TaqMan PCR Master Mix (Life Technologies) using the 7500 Fast real-time PCR system (Life Technologies). cDNA samples were loaded in duplicates, and the expression assays used for the markers of in addition to the endogenous controls. An NTC (no-template control) reaction and a RT-ve control reaction were used to check on for unspecific amplification and amplification from gDNA respectively. Relative fold changes were calculated by using the comparative CT method ($2^{-\Delta\Delta CT}$).

### SUPPLEMENTARY TABLES

**Table S1.** Clinicopathological information of patients with depression and control subjects

**Table S1. Clinicopathological information of patients with depression and control subjects**

| Sex | Age | Race | Cause of death | PMI | pH Value | Axis 1 | Axis 1 dependence | Substance at death | Psychiatric medication last 3 months |
|---|---|---|---|---|---|---|---|---|---|
| Male | 63 | Caucasian | Accident | 13 | 6.84 | Nil | Nil | Nil | Yes |
| Male | 81 | Caucasian | Accident | 98.75 | 6.80 | Nil | Nil | AD (SSRI), AP | Yes |
| Male | 41 | Caucasian | Natural | 24 | 6.00 | Nil | Nil | Nil | Nil |
| Male | 46 | Caucasian | Natural | 19.5 | 6.42 | Nil | Nil | Nil | Nil |
| Male | 64 | Caucasian | Natural | 70 | 5.65 | Nil | Nil | Nil | N/A |
| Male | 71 | Caucasian | Natural | 17 | 6.20 | Nil | Nil | Nil | Nil |
| Male | 43 | Caucasian | Natural | 27 | 6.70 | Nil | Nil | Cbd + Metab | Nil |
| Male | 51 | Caucasian | Accident | 15 | 6.83 | Nil | Nil | Eth | Nil |
| Male | 55 | Caucasian | Accident | 24 | 6.75 | Nil | Nil | Nil | Nil |
| Male | 40 | Caucasian | Accident | 24 | 6.32 | Nil | SD | Opt, BZ | N/A |
| Male | 42 | Caucasian | Accident | 63 | 6.75 | Nil | Nil | Nil | Nil |
| Male | 59 | Caucasian | Accident | 72.78 | 6.76 | Nil | Nil | Nil | N/A |
| Male | 26 | Caucasian | Accident | 12 | 6.75 | Nil | Nil | Eth, Cbd | Nil |
| Male | 42 | Caucasian | Natural | 20 | 6.62 | Nil | Nil | BZ | BZ |
| Male | 47 | Caucasian | Natural | 12 | 6.49 | Nil | Nil | Nil | Nil |
| Male | 54 | Caucasian | Natural | 25.25 | 6.61 | Nil | Nil | N/A | N/A |
| Male | 52 | Caucasian | Natural | 72.5 | 6.11 | Nil | Nil | Nil | Nil |
| Male | 55 | Caucasian | Natural | 27.5 | 5.80 | Nil | Nil | Nil | N/A |
| Male | 48 | Caucasian | Natural | 14 | 6.25 | Nil | Nil | Eth | Nil |
| Male | 57 | Caucasian | Natural | 115.3 | 6.34 | MDD | SD | BZ, AD (TCA)+Metb, Eth | N/A |
| Male | 40 | Caucasian | Suicide | 23 | 6.21 | MDD | SD | Nil | N/A |
| Male | 42 | Caucasian | Suicide | 21 | 6.40 | MDD | Nil | AD (TCA) | Classic AD, BZ |
| Male | 45 | Caucasian | Suicide | 20.5 | 6.57 | MDD | SD | Eth | N/A |
| Male | 68 | Caucasian | Suicide | 32 | 6.93 | dD, NOS | Nil | Nil | N/A |
| Male | 67 | Caucasian | Suicide | 56 | 6.85 | MDD | Nil | Nil | Nil |
| Male | 77 | Caucasian | Suicide | 26.75 | 6.30 | dD, NOS | Nil | N/A | Nil |
| Male | 64 | Caucasian | Suicide | 27.75 | 6.25 | dD, NOS | Nil | AD (SSRI) | AD (SSRI) |
| Male | 53 | Caucasian | Suicide | 29 | 6.30 | dD, NOS | SD | Nil | N/A |
| Male | 53 | Caucasian | Suicide | 14 | 6.64 | dD, NOS | Nil | N/A | N/A |
| Male | 48 | Caucasian | Suicide | 21.5 | 6.79 | MDD | SD | Eth, AD (SSRI), BZ | AD (SSRI) |
| Male | 39 | Caucasian | Suicide | 90 | 6.74 | dD, NOS | Nil | Eth | N/A |
| Male | 40 | Caucasian | Suicide | 20 | 6.33 | MDD | SD | Eth, BZ, Coc+Metb | AD (SSRI), BZ |
| Male | 42 | Caucasian | Suicide | 64 | 6.78 | MDD | Nil | Eth, DPH+Metb | AD (SSRI) |
| Male | 52 | Caucasian | Suicide | 86.5 | 6.20 | MDD | SD | Eth, Coc | Nil |

| Sex | Age | Race | Cause of death | PMI | pH Value | Axis 1 | Axis 1 dependence | Substance at death | Psychiatric medication last 3 months |
|---|---|---|---|---|---|---|---|---|---|
| Female | 66 | Caucasian | Accident | 61 | 6.80 | Nil | Nil | N/A | Nil |
| Female | 76 | Caucasian | Accident | 26.5 | 6.50 | Nil | Nil | Nil | N/A |
| Female | 81 | Caucasian | Natural | 83 | 6.50 | Nil | Nil | Nil | BZ |
| Female | 72 | Caucasian | Natural | 17 | 6.10 | Nil | Nil | N/A | Nil |
| Female | 51 | Caucasian | Natural | 111.32 | 6.50 | Nil | Nil | AD (SSRI) | Classic AD |
| Female | 68 | Caucasian | Natural | 74.3 | 6.21 | Nil | Nil | Mor, AH | N/A |
| Female | 81 | Caucasian | Natural | 44.63 | 5.91 | Nil | Nil | BZ, DPH | Nil |
| Female | 49 | Caucasian | Accident | 67.25 | 6.81 | Nil | Nil | DPH | N/A |
| Female | 79 | Caucasian | Accident | 61.5 | 6.40 | Nil | Nil | Barb, BZ | Nil |
| Female | 82 | Caucasian | Natural | 106 | 7.00 | Nil | Nil | Nil | N/A |
| Female | 40 | Caucasian | Natural | 106.5 | 6.50 | dD, NOS | Nil | N/A | Nil |
| Female | 65 | Caucasian | Suicide | 64 | 6.31 | MDD | Nil | BZ, Cd | AD (SSRI, SARI, SNDRI & NaSSA), BZ |
| Female | 49 | Caucasian | Suicide | 59.5 | 7.50 | MDD | Nil | Opd, BZ, Opt | AD (SSRI), BZ |
| Female | 55 | Caucasian | Suicide | 26.25 | 6.50 | MDD | Nil | Eth | Antimanic |
| Female | 75 | Caucasian | Suicide | 97 | 6.50 | MDD | Nil | AD (SNRI), BZ | AD (SNRI), AP |
| Female | 85 | Caucasian | Suicide | 87 | 6.50 | MDD | Nil | ßB, AD (SSRI) | AD (SSRI & SNRI) |
| Female | 80 | Caucasian | Suicide | 46.9 | 7.00 | dD, NOS | Nil | AD (SSRI) and metab | AD (SSRI) |
| Female | 59 | Caucasian | Suicide | 25.6 | 6.27 | MDD | Nil | AD (NaSSA) | AD (SSRI), BZ |
| Female | 25 | Caucasian | Suicide | 20 | 6.73 | dD, NOS | Nil | Nil | N/A |
| Female | 46 | Caucasian | Suicide | 15 | 6.53 | MDD | Nil | Eth, Barb, BZ | AD (SSRI & NaSSA), AP, BZ |
| Female | 40 | N/D | Suicide | 49.5 | 6.81 | MDD | Nil | AC, AD (SNRI ), BZ | AD (TCA & SNRI), BZ, AC |
| Female | 25 | Caucasian | Suicide | 56 | 6.55 | MDD | Nil | N/A | AP |
| Female | 51 | Caucasian | Suicide | 36 | 6.86 | MDD | SD | Cd, Eth, Opt | AD (SSRI), BZ |
| Female | 44 | Caucasian | Suicide | 60 | 6.86 | MDD | SD | Eth | Nil |
| Female | 32 | Caucasian | Suicide | 41 | 6.89 | MDD | Nil | N/A | AD (SSRI), BZ |
| Female | 41 | Caucasian | Suicide | 54.25 | 6.70 | MDD | Nil | Nil | N/A |
| Female | 48 | Caucasian | Suicide | 36.75 | 6.50 | MDD | Nil | BZ, AD (Non-TCA & SNRI | N/A |

**Abbreviations: AC, anticonvulsant; AD, antidepressant: AH, anti-histamine; AP, antipsychotic; Barb, barbiturate; ßB, ß-blocker; BZ, benzodiazepine; Cbd, cannabinoid; Cd, codeine; Coc, cocaine; dD, depressive disorders; DPH, diphenhydramine; Eth, ethanol; MDD, major depressive disorder; Metab, metabolite; Mor, morphine; NaSSA, noradrenergic and specific serotonergic antidepressant; N/A, not available; NOS, not otherwise specified; Opd, opioid; Opt, opiate; SARI, serotonin antagonist and reuptake inhibitor; SD, substance dependence; SNDRI, serotonin-norepinephrine-dopamine reuptake inhibitor; SNRI, serotonin and noradrenaline reuptake inhibitor; SSRI, selective serotonin reuptake inhibitor; TCA, tricyclic antidepressant**

**Table S2.** Clinicopathological information of subjects used for RNA sequencing

| Sex | Age | PMI | Cause of death | Tissue Source |
|---|---|---|---|---|
| M | 22 | 24.02 | Brain edema | NIH Neurobiobank |
| M | 28 | 7 | Multiple injuries, accident | NIH Neurobiobank |
| M | 33 | 24 | Fentanyl intoxication and ASCVD | NIH Neurobiobank |
| M | 38 | 20 | Neck injuries | NIH Neurobiobank |
| M | 38 | 10.45 | Wegener disease, vasculitis, aluminium intoxication | NBB |
| M | 50 | 8.3 | Cardiac arrest | NBB |
| M | 53 | 14.25 | Heart failure | NBB |
| M | 55 | 7.3 | Euthanasia, esophageal cancer | NBB |
| M | 57 | 4.25 | Myocardial infarction | NBB |
| M | 62 | 7.2 | N.A. | NBB |
| M | 65 | 5.12 | Heart failure | NBB |
| M | 71 | 7.4 | Sepsis, hypertensive cardiomyopathy | NBB |
| M | 74 | 8 | Myocardial infarction | NBB |
| M | 79 | 7.4 | Bronchopneumonia and sepsis | NBB |
| F | 47 | 4 | Respiratory failure, metastasized mamma carcinoma | NBB |
| F | 50 | 4.1 | Metastasized large cell bronchocarcinoma | NBB |
| F | 52 | 6.5 | Leiomyosarcoma with metastasis | NBB |
| F | 58 | 6.15 | Multiple organ failure | NBB |
| F | 60 | 6.5 | Endstage metastasized mamma carcinoma | NBB |
| F | 69 | 6.15 | Cardiogenic shock | NBB |

**Table S3**. Raw CT values for TAC, TACR1, TACR2, TACR3 and Substance P in the five regions of male and female controls and depressed suicide subjects

| | | TAC | TACR1 | TACR2 | TACR3 | SUBSTANCE P |
|---|---|---|---|---|---|---|
| PFC | Male Controls | 27.0 ± 0.6 | 31.4 ± 0.4 | 32.4 ± 0.3 | 30.9 ± 0.5 | 25.4 ± 0.2 |
| | Male Suicides | 25.8 ± 0.08 | 30.6 ± 0.1 | 31.5 ± 0.1 | 30.1 ± 0.4 | 24.7 ± 0.1 |
| | Female Controls | 27.0 ± 0.6 | 31.2 ± 0.3 | 32.05 ± 0.3 | 30.2 ± 3.2 | 25.5 ± 2.7 |
| | Female Suicides | 27.0 ± 0.5 | 31.6 ± 0.3 | 32.0 ± 0.3 | 30.3 ± 0.4 | 24.9 ± 0.1 |
| *Mean±S.E.M.* | | *26.7±0.3* | *31.2±0.2* | *32±0.2* | *30.4±0.2* | *25.1±0.4* |
| ACC | Male Controls | 27.1 ± 0.6 | 31.0 ± 0.5 | 32.4 ± 0.4 | 31.4 ± 0.6 | 26.4 ± 0.7 |
| | Male Suicides | 25.8 ± 0.7 | 30.6 ± 0.5 | 32.0 ± 5.2 | 31.1 ± 5.0 | 25.1 ± 0.4 |
| | Female Controls | 27.4 ± 0.6 | 31.3 ± 0.6 | 32.3 ± 0.4 | 31.9 ± 0.7 | 26.5 ± 0.6 |
| | Female Suicides | 26.7 ± 0.4 | 31.3 ± 0.3 | 32.6 ± 0.2 | 31.6 ± 0.4 | 26.3 ± 0.6 |
| *Mean±S.E.M.* | | *26.8±0.3* | *31.1±0.2* | *32.3±0.1* | *31.5±0.2* | *26.1±0.3* |
| DRN | Male Controls | 20.2 ± 2.01 | 26.3 ± 1.4 | 29.5 ± 0.8 | 27.68 ± 0.74 | 22.6± 1.1 |
| | Male Suicides | 20.2 ± 0.5 | 26.5 ± 0.3 | 28.8 ± 0.5 | 26.6 ± 0.4 | 20.7± 0.3 |
| | Female Controls | 19.2 ± 0.7 | 25.9 ± 0.2 | 28.7 ± 0.2 | 27.4 ± 0.6 | 21.4± 0.5 |
| | Female Suicides | 19.9 ± 0.5 | 26.7 ± 0.7 | 29.2 ± 0.5 | 25.8 ± 0.6 | 20.9± 0.5 |
| *Mean±S.E.M.* | | *19.9±0.2* | *26.4±0.2* | *29.1±0.2* | *26.9±0.4* | *21.4±0.4* |
| LC | Male Controls | 23.8 ± 1.2 | 26.6 ± 0.8 | 29.6 ± 0.4 | 29.9 ± 0.6 | 24.5 ± 0.8 |
| | Male Suicides | 23.3 ± 1.6 | 30.5 ± 2.3 | 32.2 ± 1.7 | 28.4 ± 0.8 | 23.3 ± 0.7 |
| | Female Controls | 24.3 ± 1.2 | 28.9 ± 1.8 | 32.9 ± 1.3 | 27.9 ± 0.5 | 22.5 ± 0.8 |
| | Female Suicides | 21.9 ± 1.6 | 29.7 ± 2.8 | 33.4 ± 1.8 | 26.7 ± 0.2 | 21.1 ± 0.3 |
| *Mean±S.E.M.* | | *23.3±0.5* | *28.9±0.8* | *32±0.8* | *28.2±0.7* | *22.9±0.7* |
| MRN | Male Controls | 25.8 ± 0.3 | 31.0 ± 0.3 | 33.9 ± 0.32 | 28.9 ± 0.4 | 25.9 ± 0.3 |
| | Male Suicides | 25.2 ± 0.4 | 30.2 ± 0.3 | 33.5 ± 0.2 | 28.3 ± 0.3 | 24.9 ± 0.5 |
| | Female Controls | 25.2 ± 0.7 | 31.0 ± 0.2 | 33.8 ± 0.2 | 29.1 ± 0.4 | 25.5 ± 0.6 |
| | Female Suicides | 25.9 ± 1.1 | 31.2 ± 0.9 | 33.8 ± 0.3 | 29.5 ± 0.8 | 27.0 ± 0.8 |
| *Mean±S.E.M.* | | *25.5±0.2* | *30.9±0.2* | *33.8±0.1* | *29±0.3* | *25.8±0.4* |

**Table S4. Raw CT values for NPY, NPYR1, NPYR2 and NPYR5 in the five regions of male and female controls and depressed suicide subjects**

| | | NPY | NPYR1 | NPYR2 | NPYR5 |
|---|---|---|---|---|---|
| PFC | Male Controls | 19.47±3.77 | 21.92±3.99 | 26.17±4.09 | 26.77±2.02 |
| | Male Suicides | 19.09±3.58 | 20.76±1.89 | 26.99±2.09 | 26.49±1.86 |
| | Female Controls | 19.64±2.08 | 21.73±2.96 | 26.85±2.78 | 26.76±1.43 |
| | Female Suicides | 18.30±2.2 | 20.21±1.92 | 26.34±1.54 | 25.91±1.06 |
| *Mean±S.E.M.* | | *19.13±0.30* | *21.15±0.40* | *26.59±0.20* | *26.48±0.20* |
| ACC | Male Controls | 19.96±0.65 | 23.01±0.36 | 29.18±0.22 | 26.98±0.35 |
| | Male Suicides | 20.06±0.52 | 23.26±0.37 | 29.34±0.47 | 26.4±0.19 |
| | Female Controls | 20.66±0.61 | 23.24±0.50 | 29.02±.29 | 26.76±0.14 |
| | Female Suicides | 19.63±0.64 | 23.07±0.44 | 29.83±0.26 | 26.88±0.18 |
| *Mean±S.E.M.* | | *20.08±0.21* | *23.14±0.06* | *29.34±0.18* | *26.78±0.11* |
| DRN | Male Controls | 26.19±1.75 | 26.97±1.6 | 27.27±1.43 | 27.81±1.84 |
| | Male Suicides | 25.98±2.01 | 26.80±2.5 | 27.20±2.20 | 27.75±1.58 |
| | Female Controls | 25.84±1.34 | 26.78±2.17 | 26.70±1.41 | 27.65±1.77 |
| | Female Suicides | 25.74±2.72 | 25.35±3.43 | 26.23±2.12 | 27.86±0.96 |
| *Mean±S.E.M.* | | *25.94±0.10* | *26.47±0.38* | *26.85±0.24* | *27.74±0.06* |
| LC | Male Controls | 26.58±0.34 | 25.01±0.49 | 26.56±0.62 | 26.76±0.33 |
| | Male Suicides | 24.96±0.22 | 24.48±0.24 | 29.15±0.80 | 27.75±0.69 |
| | Female Controls | 23.76±0.92 | 25.26±0.51 | 27.02±0.41 | 26.55±0.22 |
| | Female Suicides | 22.83±1.20 | 24.92±0.66 | 28.27±0.63 | 27.32±0.37 |
| *Mean±S.E.M.* | | *24.53±0.81* | *24.92±0.16* | *27.75±0.59* | *27.10±0.27* |
| MRN | Male Controls | 26.88±3.05 | 30.01±2.52 | 28.84±0.96 | 31.24±1.57 |
| | Male Suicides | 26.67±1.76 | 29.68±2.34 | 29.09±1.80 | 32.01±2.50 |
| | Female Controls | 27.05±1.08 | 30.29±2.67 | 28.80±2.08 | 31.84±1.92 |
| | Female Suicides | 27.26±1.83 | 31.28±1.62 | 28.98±2.17 | 31.64±2.03 |
| *Mean±S.E.M.* | | *26.99±0.15* | *30.32±0.34* | *28.93±0.07* | *31.68±0.17* |

**Table S5.** Raw CT values for Gal, GalR1, GalR2 and GalR3 in the five regions of male and female controls and depressed suicide subjects

| | | Gal | GalR1 | GalR2 | GalR3 |
|---|---|---|---|---|---|
| PFC | Male Controls | 27.64±0.42 | 26.19±0.21 | 31.96±0.27 | 33.14±0.23 |
| | Male Suicides | 27.07±0.25 | 25.24±0.3 | 30.93±0.22 | 32.97±0.17 |
| | Female Controls | 27.38±0.33 | 25.92±0.35 | 31.45±0.16 | 32.9±0.17 |
| | Female Suicides | 26.69±0.24 | 25.50±0.26 | 31.43±0.21 | 33.07±0.25 |
| *Mean±S.E.M.* | | *27.19±0.2* | *25.71±0.21* | *31.44±0.21* | *33.02±0.05* |
| ACC | Male Controls | 27.52±0.44 | 28.66±0.3 | 31.64±0.32 | 33.9±0.31 |
| | Male Suicides | 27.59±0.35 | 28.49±0.36 | 31.32±0.29 | 33.49±0.25 |
| | Female Controls | 27.30±1.38 | 28.7±0.49 | 31.66±0.42 | 33.58±0.3 |
| | Female Suicides | 27.74±0.35 | 28.5±0.31 | 32.05±0.23 | 32.92±0.13 |
| *Mean±S.E.M.* | | *27.54±0.09* | *28.59±0.05* | *31.67±0.15* | *33.47±0.2* |
| DRN | Male Controls | 27.46±0.4 | 28.1±0.23 | 28.86±0.34 | 32.19±0.3 |
| | Male Suicides | 25.59±0.14 | 26.3±0.26 | 28.9±0.18 | 30.73±0.31 |
| | Female Controls | 26.35±0.32 | 27.19±0.38 | 29.4±0.14 | 32.16±0.24 |
| | Female Suicides | 25.58±0.1 | 26.5±0.29 | 29.88±0.29 | 30.96±0.23 |
| *Mean±S.E.M.* | | *26.25±0.44* | *27.02±0.41* | *29.26±0.24* | *31.51±0.39* |
| LC | Male Controls | 26.04±0.25 | 27.97±0.48 | 32.63±0.2 | 31.88±0.27 |
| | Male Suicides | 24.24±0.17 | 25.43±0.2 | 32.10±0.24 | 30.13±0.19 |
| | Female Controls | 24.44±0.4 | 26.57±0.3 | 32.47±0.36 | 31.89±0.12 |
| | Female Suicides | 23.79±0.38 | 25.85±0.3 | 32.12±0.28 | 30.54±0.36 |
| *Mean±S.E.M.* | | *24.63±0.49* | *26.45±0.56* | *32.33±0.13* | *31.36±0.63* |
| MRN | Male Controls | 25.25±0.18 | 26.79±0.29 | 34.65±0.27 | 32.95±0.26 |
| | Male Suicides | 24.04±0.15 | 25.8±0.23 | 35.37±0.21 | 31.89±0.21 |
| | Female Controls | 26±0.25 | 27.53±0.28 | 34.36±0.27 | 32.32±0.13 |
| | Female Suicides | 25.29±0.56 | 27.59±0.74 | 34.27±0.33 | 30.67±0.5 |
| *Mean±S.E.M.* | | *25.15±0.41* | *26.93±0.42* | *34.66±0.25* | *31.96±0.48* |
